# On the complexity of haplotyping a microbial community

**DOI:** 10.1101/2020.08.10.244848

**Authors:** Samuel M. Nicholls, Wayne Aubrey, Kurt De Grave, Leander Schietgat, Christopher J. Creevey, Amanda Clare

**Affiliations:** Department of Computer Science, Aberystwyth University, Aberystwyth, UK; Department of Computer Science, Katholieke Universiteit Leuven, Leuven, Belgium; Institute of Biological, Rural and Environmental Sciences, Aberystwyth University, Aberystwyth, UK; Flanders Make, Lommel, Belgium; Institute of Global Food Security, School of Biological Sciences, Queen’s University, Belfast, UK; Institute of Microbiology and Infection, School of Biosciences, University of Birmingham, Birmingham, UK

## Abstract

**Motivation:** Population-level genetic variation enables competitiveness and niche specialization in microbial communities. Despite the difficulty in culturing many microbes from an environment, we can still study these communities by isolating and sequencing DNA directly from an environment (metagenomics). Recovering the genomic sequences of all isoforms of a given gene across all organisms in a metagenomic sample would aid evolutionary and ecological insights into microbial ecosystems with potential benefits for medicine and biotechnology. A significant obstacle to this goal arises from the lack of a computationally tractable solution that can recover these sequences from sequenced read fragments. This poses a problem analogous to reconstructing the two sequences that make up the genome of a diploid organism (*i*.*e*. haplotypes), but for an unknown number of individuals.

**Results:** The problem of single individual haplotyping (SIH) was first formalised by Lancia *et al* in 2001. Now, nearly two decades later, we discuss the complexity of “haplotyping” metagenomic samples, with a new formalisation of Lancia *et al* ‘s data structure that allows us to effectively extend the single individual haplotype problem to microbial communities. This work describes and formalizes the problem of recovering genes (and other genomic subsequences) from all individuals within a complex community sample: which we term the metagenomic individual haplotyping (MIH) problem. We also provide software implementations of our proposed pairwise single nucleotide variant (SNV) co-occurrence matrix and greedy graph traversal algorithm.

**Availability and implementation:** Our reference implementation of the described pairwise SNV matrix (`Hansel`) and greedy haplotype path traversal algorithm (`Gretel`) are open source, MIT licensed and freely available online at github.com/samstudio8/hansel and github.com/samstudio8/gretel, respectively.

**Contact**

## Introduction

The problem of single individual haplotyping (SIH) was first described by Lancia *et al* at Celera Genomics in 2001 [1]. In the wake of the announcement of Celera’s first human genome, it became clear that the next big research problem was not only to analyse the millions of single point variants that populate our genomes, but how to assemble the two haplotypes that make up a single individual’s genome. This 2001 work introduced the first terminology and notation for “computational SNPology”, a phrase that did not seem to catch on in the literature. A full discussion of the history of human (and by extension, diploid) haplotyping is outside the scope of this work (although for an expanded discussion see [2]), but it is important to introduce this first description of the problem, as it formed the foundation for many other approaches and algorithms that followed.

The computational problems involved with haplotyping arise because genomic assembly algorithms typically achieve accuracy through the availability of high sequencing coverage. Although it is now possible to assemble high quality, contiguous sequences of entire genomes, even for a complex community sample [3], such sequences only represent a consensus of the true haplotypes that exist in a sample. This is problematic for the study of metagenomes where the objective of consensus generating assembly methods is at odds with our desire to explore the individual haplotypes that provide the population-level diversity of natural microbiomes. Specialized metagenomic assemblers like `flye` [4] and `metaSPAdes` [5] do not aim to reconstruct haplotypes from a microbial community. Not only is it technically challenging to sequence every true haplotype to a depth sufficient for confident assembly, but even high-quality, single-molecule, long-read sequencing platforms fail to achieve perfect recall and perfect precision on the single read level, and are therefore not capable of single-molecule haplotype identification [6]. Even high-accuracy sequencing techniques such as circular consensus sequencing still produce a per-base error rate of around 1% (and a higher error rate for insertions and deletions) that will be indistinguishable from low frequency haplotypes [7]. Ideally large-scale culturing projects would aim to culture and sequence every genome from every individual in a microbial community. However such endeavours would be incredibly laborious and even large international projects such as the Hungate Collection [8] and Human Gastrointestinal Bacteria Culture Collection [9] generally produce a subset of representative genomes for each species. Despite these difficulties, in order to reconstruct haplotypes, we must first define the problem, then work to find tractable solutions. Lancia *et al* ‘s work first described the problem *for a diploid individual* as:

> “Given a set of fragments obtained by DNA sequencing from the two copies of a chromosome, reconstruct two haplotypes that would be compatible with all the fragments observed.” — Lancia *et al*. (2001)

It is important to note that even in an ideal scenario with error-free read fragments, lack of coverage across the haplotypes (be it a chromosome, or region of interest) necessitates a problem definition where at best, one can only recover the two haplotypes that are most “compatible” with the observed fragments. Perhaps more importantly for the context of this work, Lancia also defined a common notation to formally describe the problem of single individual haplotyping. The “SNP matrix” (typically denoted *M*) is an *m* × *n* matrix encoding the binary allele observed at each SNP site 1..*n* on each read fragment 1..*m*. That is, *M* [*i*][*j*] is one of two possible alleles (typically labelled 0, 1 or *A, B*), or a gap (denoted −) observed at the *j*’th SNP site on the *i*’th read fragment (Figure 1).

**Figure 1:**
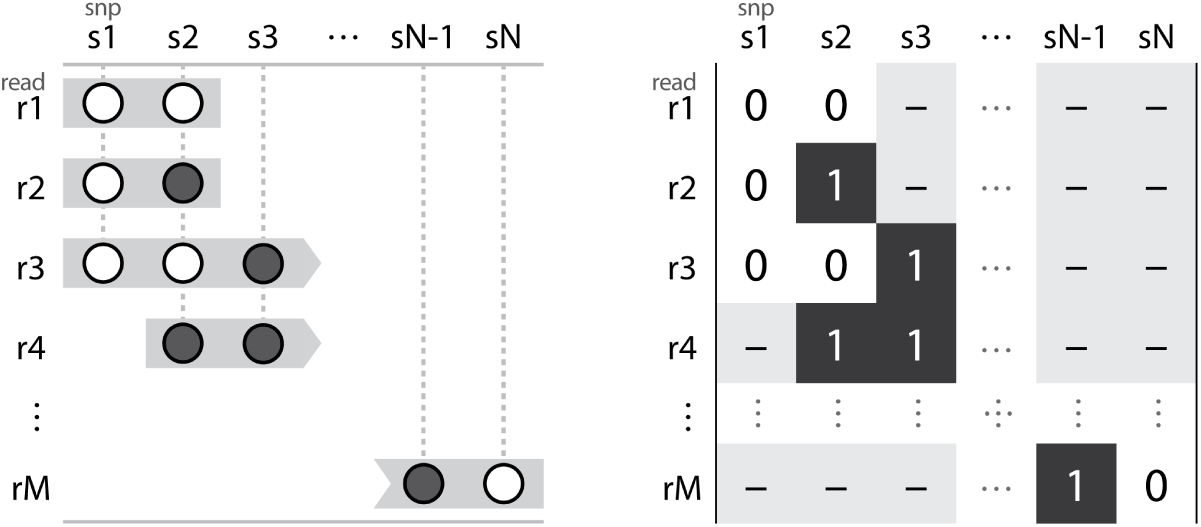
An example SNP matrix. Read fragments represented by grey boxes (left) are aligned to some reference with known SNP loci. The alleles at the SNP loci are represented by white and grey circles. These reads can be alternatively represented by an *m* × *n* SNP matrix (right). Each row of the matrix models one of the *m* read fragments and each column corresponds to one of the *n* SNPs. Elements encode the allele at a given SNP for a particular read fragment as a 0 or 1, or a − if the read does not cover that position. A column containing only one element indicates the corresponding SNP site is homozygous, otherwise it is heterozygous.

Although unstated, it would appear the inspiration for Lancia’s SNP matrix in 2001 is likely from a data structure introduced by Churchill and Waterman in 1992 [10]. Churchill and Waterman described a matrix with *m* read fragment rows and *n* base position columns. An element in this matrix (which given the benefit of hindsight, we will denote as *M*), *M* [*i, j*] refers to the nucleotide observed “on a gel” at position *j* on the *i*’th read fragment. Of course, the problem a decade prior considers the accuracy of only one sequence: that of the true DNA molecule that was sequenced, rather than untangling diploid sequence data to recover a solution of two distinct haplotypes. Lancia *et al* ‘s work additionally defines three optimisation problems to solve SIH, which involve identifying and mitigating different types of “conflicts” in the SNP matrix. A pair of read fragments *r*_*i*_, *r*_*j*_ are said to be in fragment conflict if they have opposing alleles on at least one SNP. Similarly, a pair of SNPs *s*_*k*_, *s*_*l*_ are said to be in SNP conflict if reads *r*_*u*_ and *r*_*v*_ are heterozygous at one SNP and homozygous at the other. Although a conflict might indicate that a pair of fragments originate from different haplotypes, Lancia *et al*. focus on the idea that conflicts arise due to errors in the underlying sequence data, arguing that “experiments in molecular biology are never error-free”. A SNP matrix *M* which contains any conflicts is *infeasible*, and the three optimisation methods aim to graphically represent and resolve the conflicts in *M*, to yield a pair of feasible haplotypes. This SNP matrix influenced almost all haplotyping algorithms for the next twenty years [2, 11].

We take inspiration from this seminal work and offer a new data structure and algorithmic solution to generalise the haplotype recovery problem to individuals in a microbial community. We term this the metagenome individual haplotyping (MIH) problem.

## Theory

We begin with a naive *de novo* formulation of the metagenomic individual haplotyping problem (MIH), where the input is a collection of reads generated by a DNA sequencer from an environmental sample, with an unknown number of organisms. The ideal output from a solution to the MIH problem is the collection of whole-genome sequences representing all the individual organisms in a microbial community. To illustrate further, we require the following definitions:

- Ω, a microbial community.
- *O*, a set containing each full genomic sequence, of each individual organism in environment Ω. *O* is *the* “metagenome” of Ω: encompassing all possible genomes in the environment.

Arguably one could consider *O* as a bag, allowing multiple copies of the exact same genome in the community. For the purposes of haplotyping we do not need to concern ourselves with recovering duplicate genomes. We also require some definitions to denote how the community is sampled:

- *σ* = Sample(Ω)
  - A sample taken from microbial community Ω.
  - *σ* ⊆ Ω
- *M*
  - A set containing the unique genomes *m* ∈ *M*, from the individual organisms captured in the sample *σ*.
  - *M* represents the metagenome that was captured in the sample *σ*, and is our insight into *O*.
  - As a subsample, *M* is not necessarily representative of the entire metagenome *O*.
  - *M* ⊆ *O*
- *R* = Sequencing(*σ*)
  - *R* is the set of **reads** obtained from the sequencing of isolated DNA from sample *σ*.
  - A read *m*[*u*: *v*] describes a fragment of some genome *m* ∈ *M*, covering positions *u, v* ∈ 1..|*m*|, with some degree of error.
  - Additionally, due to library preparation bias and sequencing errors, *R* is unlikely to provide uniform and non-zero coverage of the bases across all *m* ∈ *M*.

Formally, the goal of MIH appears to be the recovery of *M* from *R*. However, to construct a form of SNP matrix, we must identify the variants that comprise haplotypes, which in turn necessitates a common frame of reference for the reads. In the context of SIH, this would be a known reference sequence. Unfortunately for us, the ideal frame of reference for MIH is *M* - the very thing we are trying to recover. Thus, our naive definition of *de novo* MIH collapses to an exhaustive, special case of the *de novo* assembly problem. A solution to MIH is confounded by five problems: (a) DNA from every genome needs to be extracted and sequenced to a depth sufficient for recovery, (b) genomes share homologous regions that require disambiguation, (c) reads may be of an insufficient length to disambiguate repeats or resolve bridges between variants, (d) sequencing error can be indistinguishable from rare haplotypes and (e) the presence of an unknown number of haplotypes complicates the already computationally difficult (NP-hard) [12] problem of haplotyping. These issues all apply to both the metagenomic haplotyping and metagenomic assembly problems, which is why there is no metagenomic *de novo* assembler that attempts to exactly recover *M* from *R*. Metagenomic *de novo* assemblers such as `flye` and `metaSPAdes` still aim to recover consensus sequences. Although recent experimental developments to `flye` allow a user to retain “haplotigs”, whereby bubbles in the graph structure are left uncollapsed, this still requires significant coverage for those alternative sequences to be assembled in the first place, and comes at the cost of assembly contiguity.

Although such *de novo* consensus sequences do not represent the entire metagenome of *M*, and typically collapse regions with shared homology into single broken contigs, we suggest that they provide a suitable approximation against which to align reads and construct a SNP matrix for the purpose of haplotyping. We therefore redefine the input to the MIH problem as a collection of sequencing reads *R*, their alignment to a set of *de novo* assembled contigs (and/or any existing references) and the variants determined by inspecting that alignment:

- *C* = Assemble(*R*)
  - Contig set *C* (assembly) constructed *de novo* from the reads *R* by some `Assemble` operation.
  - *c*_*k*_[*j*] ∈ {*A, C, G, T, N*} for *k* ∈ 1..|*C*|, *j* ∈ 1..|*c*_*i*_|.
  - `Assemble` attempts to construct a set of consensus sequences representative of genomes in *M*, from the reads *R*. It should be noted that `Assemble` does not try to recover *M*, and typically fails to distinguish between similar sequences that should create distinct *c* ∈ *C*.
- *A* = Align(*R, C*)
  - Alignment *A* is generated by aligning read set *R* to contig set *C* with operation `Align`.
  - An alignment *a* ∈ *A* defines a correspondence between some subsequence of read *r* ∈ *R* and a subsequence of contig *c* ∈ *C*. Note that the length of the subsequences with correspondence between *r* and *c* may not necessarily be equal.
  - 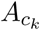 is the set of all alignments to contig *c*_*k*_.
  - 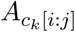 is the subset of alignments to contig *c*_*k*_ where any correspondence in *R* was found between positions *i* and *j* on contig *c*_*k*_.
- 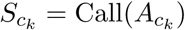
  - The list of positions on contig *c*_*k*_ ∈ *C* determined to be heterogeneous by the operation `Call`.
  - We will henceforth refer to these positions as SNVs rather than SNPs in our descriptions to make it clear that there are no assumptions on variant frequency or validity.
  - `Call` may simply consider each ‘column’ 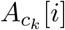 for *i* ∈ 1..|*c*_*k*_| and determine position *i* as a variant if there is a disagreement on the nucleotide at that position across the aligned reads.
- `Call` may also be a more complex variant prediction algorithm.

To enable computational tractability and reduce the reliance on the quality of the assembly, we can consider the local MIH problem. The input to the local MIH problem is constrained by selecting a contig from *C* and filtering for alignments in *A* between positions *i* and *j*:

- *c*_*k*_[*i:j*], a **region** of contig *c*_*k*_, identified as biologically interesting
- 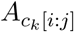, the **alignments** of reads *R* that map to the contig region *c*_*k*_[*i*: *j*]
- 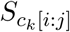, the list of positions determined to be **variants** over the region *c*_*k*_[*i*: *j*]

As *c*_*k*_[*i*: *j*] was assembled from *R*, it is an approximation of one (or more) genomes in the sampled metagenome *M*. In the same way that an assembler can collapse multiple genomes with shared sequence into single contigs, or several broken contigs, an aligner may be unable to determine whether read *r* truly belongs to a single assembled contig in *C*. We cannot guarantee that reads aligned to *c*_*k*_[*i*: *j*] originate from the same genome *m* in the sampled metagenome *M*.

We use this to our advantage for local MIH. Consider first:

- *f*, a biologically interesting genomic **feature** such as (but not limited to) a gene or operon
- *M*_*f*_, the subset of the genomes in the sampled metagenome M that have feature *f*
- *m*_*k*_, some genome in *M*_*f*_
- *m*_*k*_[*i*: *j*], a region *i*..*j* on *m*_*k*_ that has concordance with feature *f*, with appropriate length *j*−*i* ≈ |*f* |
- Γ_*f*_ = *set*(⎱*m*_*k*_[*i*: *j*] | *k* ∈ 1..|*M*_*f*_ |, *i, j* ∈ 1..|*m*_*k*_|⎰). Γ_*f*_ is the set of haplotypes of *f* in *M*.

Now let *c*_*k*_[*i*: *j*] have concordance to a feature *f*. The output of local MIH is the set of haplotypes Γ_*f*_. Although the ambiguity in the alignments 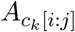 enable the variation in Γ_*f*_ to be captured, the evidence they provide does not readily resolve to haplotypes. The Lancia *et al* SNP matrix is not suitable for MIH, as the only information stored about a variant site on a read is whether it belongs to one of two haplotypes. In order to recover haplotypes that actually exist, we must define a new data structure that is capable of considering evidence from non-adjacent variant sites, and is agnostic to the number of potential haplotypes in Γ_*f*_.

## Data structure

### The pairwise SNV co-occurrence matrix

We present a new form of SNP matrix: the pairwise SNV co-occurrence matrix, denoted *H*. Consider two positions *i* and *j*, on an assembled contig *c*_*k*_. With a read *r* aligned to this contig, let *α* be the nucleotide on *r* at *i*, and *β* be the nucleotide at *j*. We say that the read supports an observation that symbol *α*_*i*_ co-occurs with symbol *β*_*j*_. *H* is a rank 4 tensor (which we will refer to as a four dimensional matrix for convenience) such that an individual element *H*[*α, β, i, j*] records the number of such observations supporting a co-occurring pair of symbols (*α*_*i*_, *β*_*j*_). Note that the reads are aligned with respect to the contig as a reference such that *α, β* are in the alphabet Σ = {*A, C, G, T, N*, −}, where − denotes a deletion in the read. We do not consider insertions.

This representation differs from the typical SNP matrix model [1] that forms the basis of many haplotyping approaches. Rather than a matrix of columns representing variants and rows representing reads, we discard the concept of a read entirely and aggregate the evidence seen across all reads with pairs of symbols and positions.

More importantly, *H* can be exploited to build other structures.

### H as a simple graph

Consider *H*[*α, β*, 1, 2] for all symbol pairs (*α, β*). One may enumerate the available transitions from position 1 to position 2. Extending this to consider *H*[*α, β, i, i* + 1] for all (*α, β*) and *i* ∈ 1..*n* − 1 (where 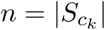 is the number of SNVs identified on contig *c*_*k*_), one can construct a simple graph *G* of possible transitions between all symbols. *G* would represent a graph of transitions observed between SNVs, across all reads. A node in *G* represents a state, pairing a symbol to a position (e.g. an ‘A’ at position 1). An edge between a pair of nodes in *G* represents two adjacent SNVs (*i, i* + 1) and can be labelled with a weight using the number of corresponding co-occurrence observations in *H*. Figure 2 shows how *H* records information about SNV pairs, and how a simple graph can be derived from this information.

**Figure 2:**
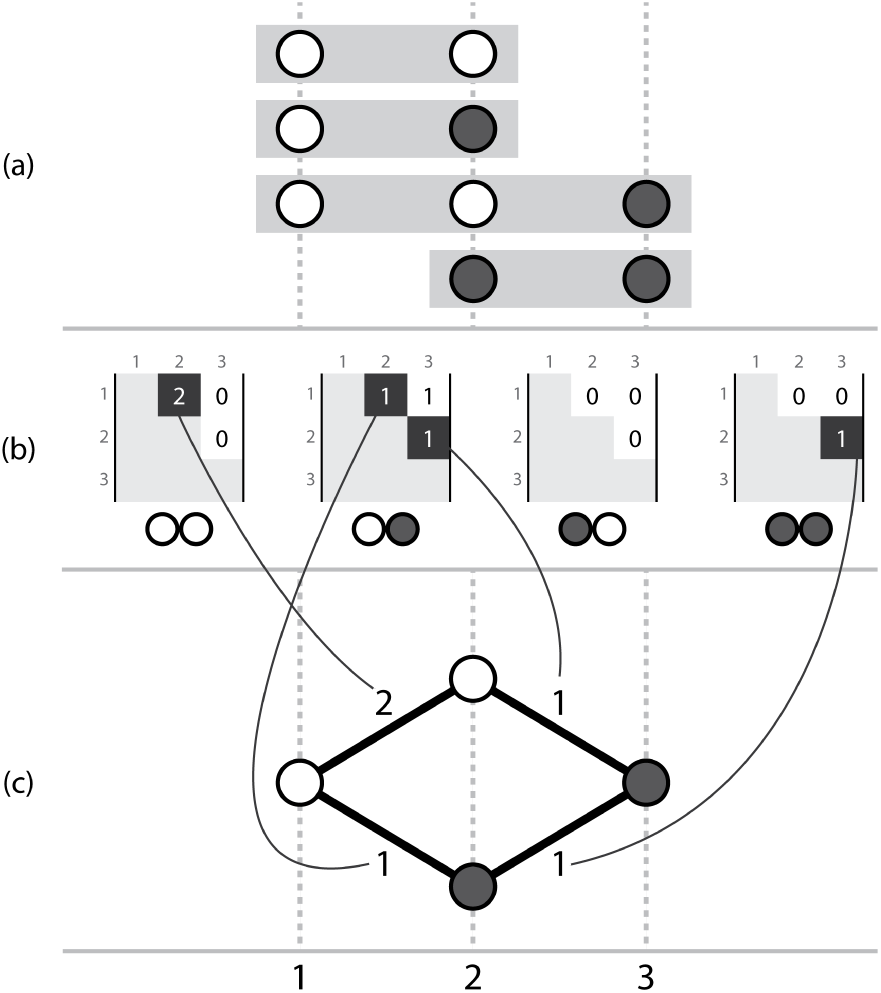
Three corresponding representations, (a) a set of aligned reads, with called variants, (b) the pairwise SNV co-occurrence matrix *H* where each possible pair of symbols (00, 01, 10, 11) has a matrix storing counts of occurrences of that ordered symbol pair between two genomic positions across the aligned reads, (c) a simple graph that can be constructed by considering the evidence provided by adjacent variants. Note for simplicity these pictorial examples only use an alphabet of two symbols.

Formally, *H* can be considered as a graph *G* = (*V, E*). Here, we define *E* and *V* as:

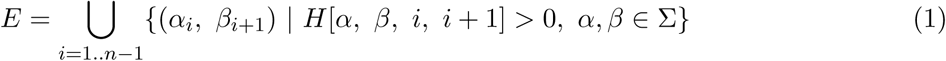

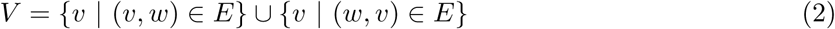

Intuitively, one may traverse a path through *G* by selecting the edges most highly supported by the corresponding evidence in *H*, in order to recover a series of symbols representing an ordered sequence of SNVs that potentially constitutes a haplotype. We can now formalize a haplotype *ĥ* as a sequence of nodes (*v* ∈ *V*) encountered in this graph:

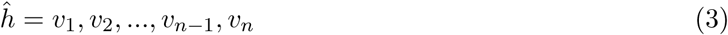

Although the analogy to a graph helps us to consider paths, the available data in *H* cannot be fully represented with a graph such as that seen in Figure 2 because it includes data about co-occurrence of non-adjacent SNV positions. A graph representation constrains our representation as edges can only be drawn between adjacent SNVs (*i, i* + 1). Without considering information about non-adjacent SNVs, one can traverse *G* to create paths that do not exist in the observed data set, as shown in Figure 3. To prevent construction of invalid paths and recover genuine paths more accurately, a SNV matrix for MIH must be able to consider evidence observed between non-adjacent symbols when determining which edge to traverse next.

**Figure 3:**
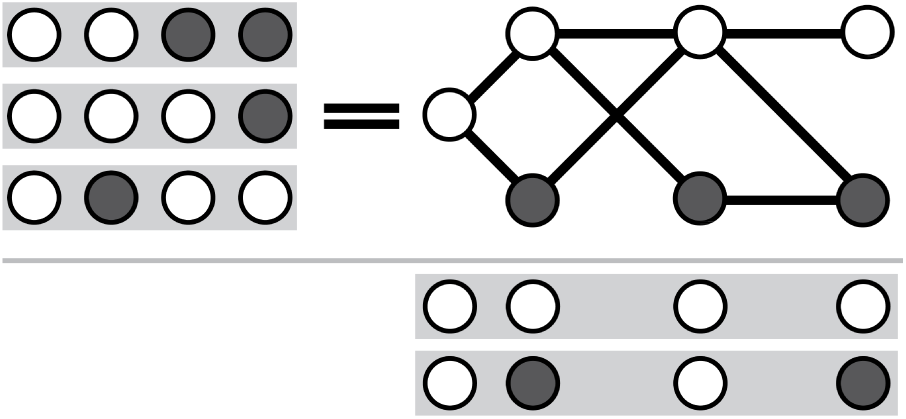
Considering only adjacent SNVs, one may create paths for which there was no actual observed evidence. Here, the reads {0011, 0001, 0100} do not support either of the results {0000, 0101}, but both are valid paths through a graph that permits edges between pairs of adjacent SNVs.

### H as a probabilistically-weighted graph

We may take advantage of the additional information available in *H* and build upon the graph *G*. Rather than directly setting the edge weights to values from *H*, we define a formula to weight edges based on the support of pairs between a path of visited nodes and any potential next node. That is, a decision to move to a symbol at position *i* + 1 is informed not only by observations in *H* supporting (*i, i* + 1), but also the non-adjacent observations (*i* − 1, *i* + 1), (*i* − 2, *i* + 1), and so on.

Here we formalize a Bayesian probabilistic framework to determine the edge weights in *G*, based on a path of already visited nodes. We define the probability of selecting *v*_*i*+1_, conditioned on the path observed so far, as:

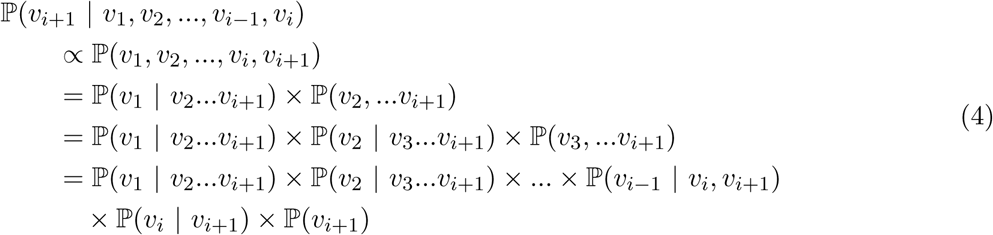

Clearly, evaluating the equation becomes more computationally expensive and risks compounding estimation errors as the path length increases. To construct a path from *v*_1_…*v*_*n*_, the upper bound for the number of operations will be |Σ|*×n* with calculations becoming increasingly complex as *i* increases. We can reduce complexity with two assumptions: (a) conditional independence between variants and (b) one can limit the number of elements in the prior path without loss of accuracy because the reads providing this co-occurrence support are limited in length. This simplifies our previous equation as we only need to consider the pairwise appearances of each *v*_*i*_ encountered thus far against *v*_*i*+1_; and limit the number of variants to consider, from the current position in the path *i*, back some small and sensible number of steps *L*:

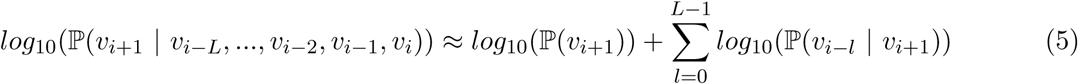

As a minor implementation detail, to overcome inaccuracies encountered through floating point error when performing mathematical operations on very small fractional values, we provide these definitions with log probabilities instead. We define *L* as the the ‘lookback’ size, the number of variants of the current path to consider when selecting *v*_*i*+1_.

We use *H* to estimate the marginal and conditional probabilities. Equation 6 provides an estimate for the marginal distribution of a symbol *β* appearing at position *j*. As a minor implementation detail, we expand our alphabet Σ to include a dummy symbol Ø. When constructing *H*, the last SNV *γ*_*j*_ on a read *r*, is automatically linked to Ø_*j*+1_ (*i*.*e. H*[*γ*, Ø, *j, j* + 1] is incremented). This allows the span between *j* and *j* + 1 to be calculated even in the case where *j* is the last observed position on a read.

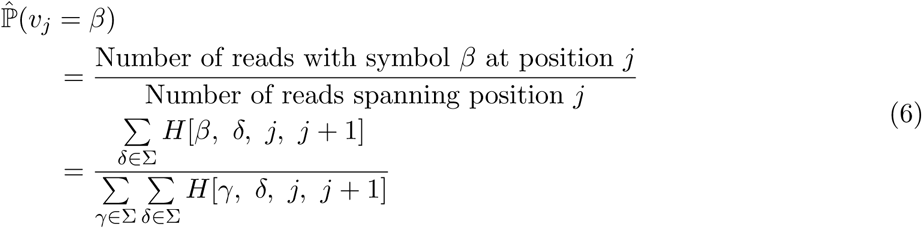

Equation 7 provides an estimate for the conditional distribution of symbol *α* appearing at position *i* given that *β* was observed at position *j*. To avoid the potential of dividing by 0 in cases where a suitable read spanning *i* and *v*_*j*_ = *β* does not exist, we apply Laplace smoothing, adding one dummy read that will provide support for each possible pair of SNV symbols (including ones that were not observed).

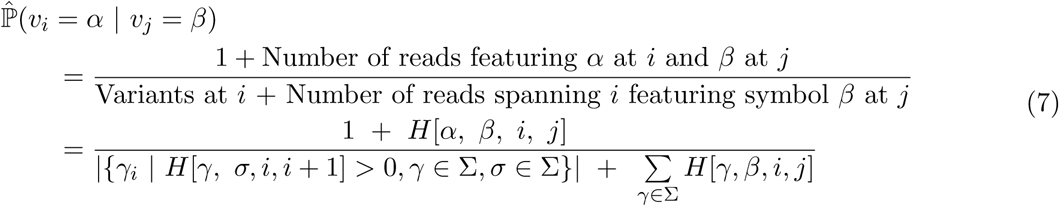

**Algorithm**

**A greedy graph traversal algorithm for local MIH**

We have formulated the above ideas into an algorithm for finding solutions to local MIH. Our algorithm traverses the graph *G* in a greedy manner, selecting the edge with the highest likelihood (given the path traversed so far) in order to recover the highest likelihood haplotypes. The core of the algorithm is as follows:

1. Given *c*_*k*_[*i*: *j*], the location of a feature of interest (*f*) in the *de novo* assembly, parse the read alignments 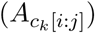 and construct the pairwise SNV co-occurrence matrix *H*
2. Initialise a haplotype *ĥ* = [], *i* = 0, and assign *L* to be the mean number of SNVs per read
3. Representing *H* as a graph *G*;
  - Query for the available edges from current position *i*, to the next position *i* + 1
  - Calculate the probabilities for each available edge using Equation 5, given *ĥ*[*i* − *L*: *i*]
  - Traverse the most likely edge and append the new node to *ĥ*, increment *i*; however if there are no edges that can be traversed, terminate the algorithm
  - If 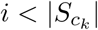 repeat step 3; else proceed to step 4
4. Report path *ĥ* as a haplotype, and modify *H* to reduce the evidence supporting *ĥ*.
5. Repeat (3-4) until encountering a node with no edges that can be traversed, or an additional stopping criterion has been reached.

### Reweighting H to find multiple haplotypes

Given *H*, the algorithm will behave deterministically and return the same haplotype for any traversal. However, the goal of local MIH is to recover Γ_*f*_: the set of haplotypes corresponding to feature *f*, rather than just one haplotype. To recover more than just one haplotype, our algorithm must modify *H* to remove evidence for that haplotype, in order to weight the calculation of edge probabilities in favour of alternative haplotypes on the next traversal. We achieve this by using the smallest marginal distribution from path *ĥ* as a ratio (*λ*) to decrease the observations in *H* that directly support the path *ĥ*. The intuition is that the least supported node in *ĥ* estimates the proportion of evidence supporting the entire haplotype.

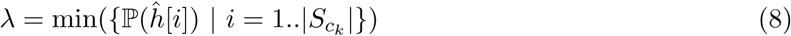

This ratio is used to decrease the evidence for all adjacent entries in *H* that correspond to *ĥ*.

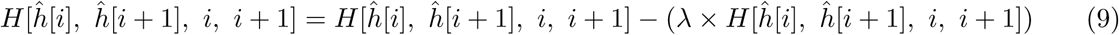

After multiple iterations of path finding and subsequent reweighting, elements in *H* will begin to approach 0, causing edges in the graph to become unavailable for traversal. At this point the algorithm will terminate. Alternatively, if this criterion is not reached after some predefined number of iterations, the algorithm will terminate.

### Using H to score and rank haplotypes

We can use the estimated probabilities to also score and rank the haplotypes recovered. For a completed haplotype, *ĥ*, we compute its likelihood based upon the sum of the marginal log probabilities for each element of *ĥ* given the state of *H* prior to reweighting.

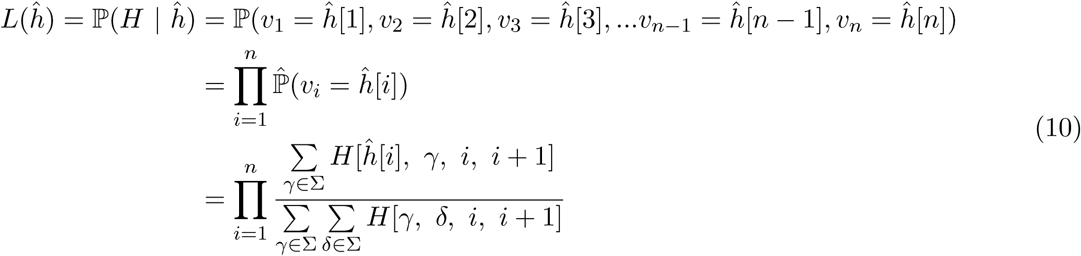

To overcome the potential for floating point arithmetic error, we calculate and report the log likelihood.

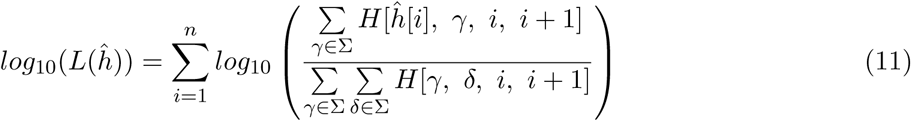

## Discussion

With recent technical advancements reducing the cost and complexity of sequencing environmental samples researchers are turning their attention to focus on the population-level variation within microbial communities. However, the field is hampered by a lack of consensus in terminology. Segata recently analysed the terminology used for metagenomic strains and for the diversity present in microbiomes [13]. He discussed the difficulties and misunderstandings that arise from a lack of strict definitions and states that it is “crucial for this line of research to at least define practical and operational definitions” to address the open problem of characterizing unknown genomes within the human microbiome. More recently, Van Rossum *et al* [14] highlighted the “overwhelming number of methods and terms to describe infraspecific variation” in a review. Their work provides much needed clarification to the terminology for metagenomic analyses. Figure 2 in their paper depicts a proposed hierarchy of terms from species, through to sub-species, strain and down to the individual genome level. Even though they have provided copious terminology definitions throughout the paper, a formalisation of the haplotype in the context of a metagenome is still absent. Our work clarifies further by providing a formal definition for the ‘metagenomic individual haplotype’.

We have defined the pairwise SNV co-occurrence matrix that packs sequencing reads into a structure compatible with metagenomic haplotyping. Our SNV matrix can be used to build a graph, reducing the problem of local MIH to that of recovering the highest weighted paths from a graph. We provide a probabilistic framework to weight the edges in this graph based on the path observed so far, while also considering evidence from non-adjacent SNVs in the underlying read data. We provide a free and open implementation of this SNV matrix (`Hansel`) at github.com/samstudio8/hansel.

Additionally, we propose a greedy solution to the problem of local MIH, using a graph representation of *H* to traverse paths through a graph to output haplotypes. Our algorithm requires no configuration, has no user-facing parameters, and requires no pre-processing of reads, other than alignment. The haplotypes recovered with our algorithm can be scored and ranked by their likelihood. We provide a free and open implementation of this algorithm (`Gretel`) at github.com/samstudio8/gretel.

Our work extends the single individual haplotyping (SIH) problem introduced by Lancia *et al*, to define the metagenomic individual haplotyping (MIH) problem. We offer our data structure and algorithm in the hope that they will form a foundation for future approaches to haplotyping microbial communities.

## Ethics approval and consent to participate

No ethical approval was necessary for this study.

## Availability of data and materials

Our reference implementation of the described pairwise SNV matrix (`Hansel`) and greedy haplotype path traversal algorithm (`Gretel`) are open source, MIT licensed and freely available online at github. com/samstudio8/hansel and github.com/samstudio8/gretel, respectively.

## Competing interests

The authors have no conflicts of interest to declare.

## Author’s contributions

All authors discussed and defined the theoretical problem. SN wrote the code and documentation. All authors contributed to the manuscript.

## Funding

For the duration of this work SN was funded via the Aberystwyth University Doctoral Career Development Scholarship and the IBERS Doctoral Programme. SN was funded to travel and work at Katholieke Universiteit Leuven by the Erasmus+ Programme. WA is funded through the Coleg Cymraeg Cenedlaethol Academic Staffing Scheme. CJC was funded by the Biotechnology and Biological Sciences Research Council (BBSRC) Institute Strategic Programme Grant, Rumen Systems Biology (BB/E/W/10964A01).

## Acknowledgements

SN wishes to acknowledge Nicholas Loman (University of Birmingham) for supporting the completion of the manuscript.

## Notes

### Competing Interest Statement

The authors have declared no competing interest.

